## Supplementary Information for "Phage-Assisted Evolution of Allosteric Protein Switches"

**The PDF file includes:**

Supplementary Note 1

Supplementary Figure 1 to 21

**Supplemental Note 1:**

Optimization of phage levels is an important factor in the successful execution of POGO-PANCE. Here we provide additional context on how input titers were chosen for each stage of the workflow and why these choices support mutation load, library diversity, and avoidance of population saturation.

Mutagenesis: Prior work (Badran & Liu, 2015) demonstrated that mutation rates for MP6 are highest when phage titers begin low and are subsequently expanded. Accordingly, each mutagenesis day was seeded at ~$10^3$ pfu/mL and expanded to $10^8$–$10^9$ pfu/mL. This approach (i) imposes a population bottleneck that favors enrichment of the most strongly selected and most abundant phage variants, and (ii) maximizes exploration of the local fitness landscape during regrowth.

For RAMPhaGE, we found that input titers remained close to output titers at high gIII induction (Supplementary Fig. 11) while maintaining high editing efficacy in the total pool. We therefore seeded initial evolutions at a high MOI (108–1010) from the outset. In this work, we performed only a single passage on Recombitron-carrying host cells prior selection, but we recommend an additional propagation step (using S2208, for example) if phage titers would be too to enter POGO-PANCE selection.

Negative Selection: Negative selection functions as a sieve that removes phages that are active under non-permissive conditions. Because each bacterium can be infected only once, the phage input must balance two competing needs. We aimed to provide enough phage to adequately sample the library generated the previous day (MOI ≤ 1), while avoiding levels that would saturate the culture (MOI > 1). This reduces the chance of allowing active phages to escape depletion and carry over into positive selection. Adjusting the IPTG concentration provides tunability in selection stringency, but it also reduces phage titer. Care should therefore be taken to avoid washout during passaging when IPTG levels are low.

Positive Selection: Negative selection removes variants that are active under the undesired condition, but it does not distinguish between variants that are correctly inactive and variants that are simply non-functional. A subsequent positive selection step is therefore essential to enrich variants that retain activity specifically under the desired condition. To ensure effective selection, we avoided saturation of the bacterial population so that productive phages could continue to infect multiple host. We thus imposed an upper boundary on the input titer to maintain consistent selective pressure (~$10^6$pfu/mL). Although this limit is not strictly required and can be adjusted according to standard PANCE principles to tune selection stringency, excessively low titers during positive selection can inadvertently seed only non-functional variants. We therefore recommend maintaining relatively high titers entering positive selection to increase the likelihood of enriching functional protein under the desired selective condition.

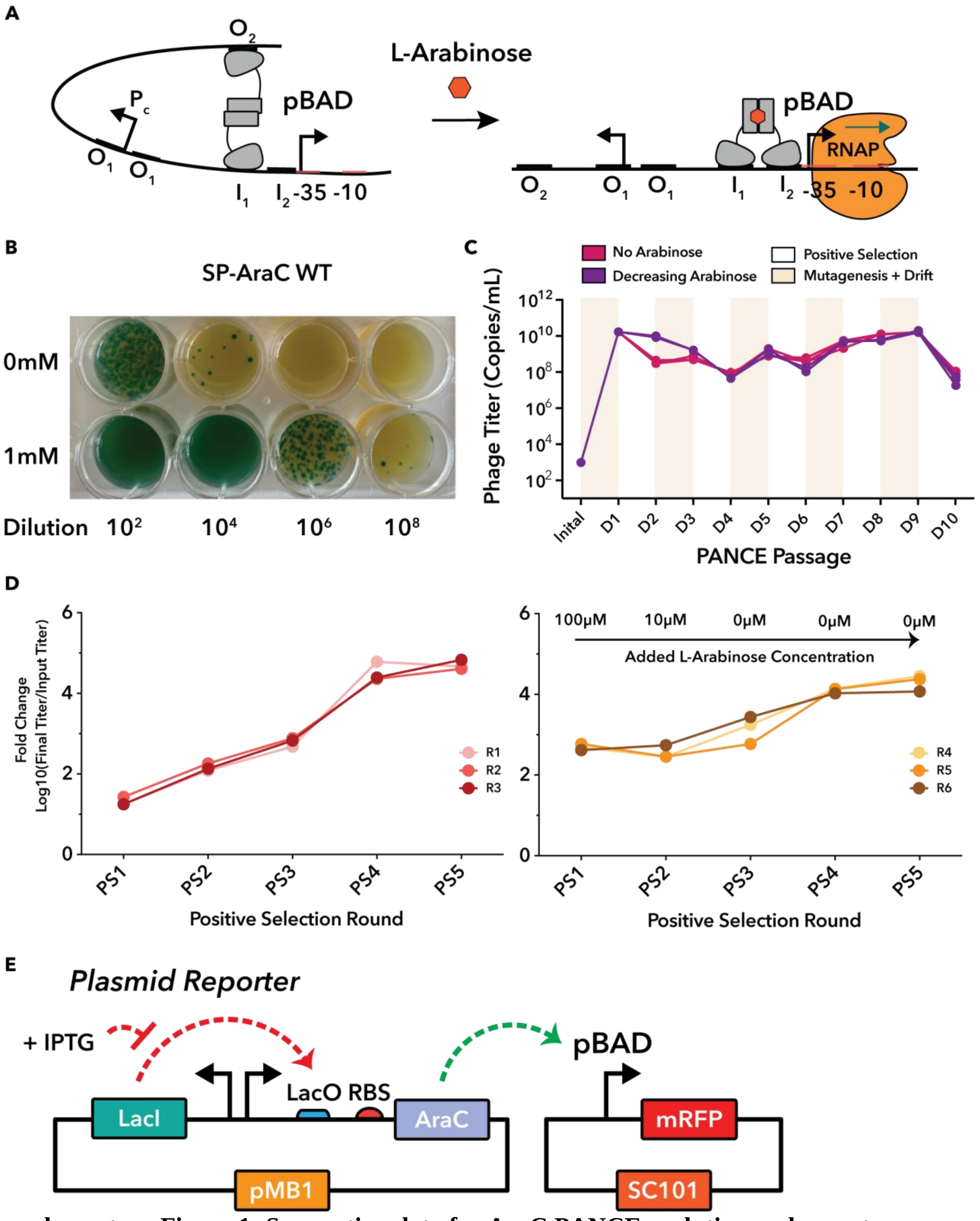


**Supplementary Figure 1: Supporting data for AraC PANCE evolution and reporter validation. a,** Schematic of AraC-mediated regulation of the pBAD promoter. In the absence of arabinose, AraC binds to distal and proximal half-sites to repress transcription. Arabinose binding shifts AraC to an activating conformation, promoting transcription initiation at pBAD. **b,** Plaque assay of phage encoding WT AraC shows arabinose-dependent propagation. **c,** qPCR quantification of phage titers across all evolution pools during the PANCE campaign. Mutagenesis and positive selection days are marked, along with pools receiving no arabinose or decreasing arabinose concentrations. **d,** Fold change in phage titer upon positive selection. Fold change was calculated by dividing the titer obtained after each positive selection day by the input dilution from the previous step, followed by $\log_{10}$ transformation. **e,** Schematic of the mRFP reporter system. In the expression plasmid, LacI suppresses expression of the AraC variant under control of a LacO-regulated promoter; IPTG addition relieves repression. The reporter plasmid encodes mRFP under the pBAD promoter.

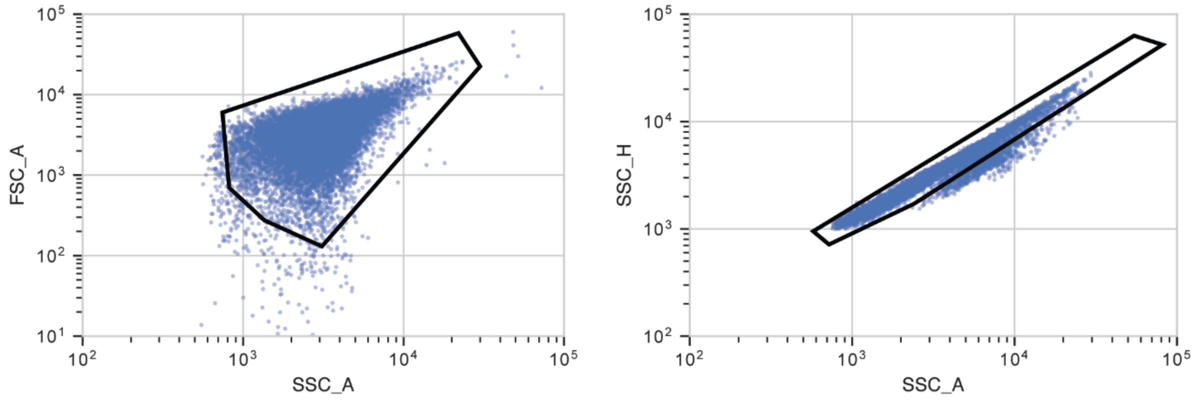


**Supplementary Figure 2: Overview of the gating strategy used to gate for single *E. coli* cells.**

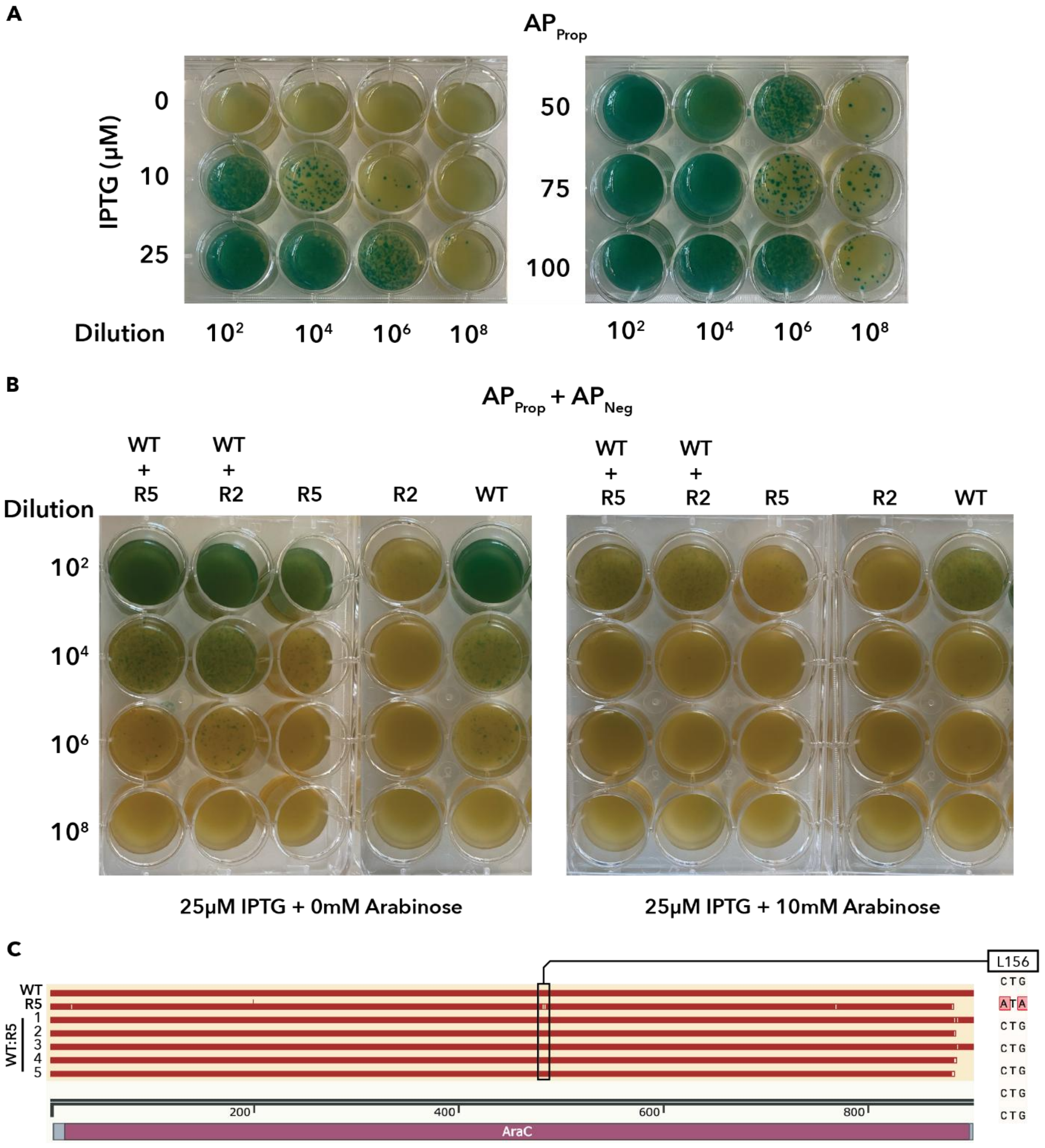


**Supplementary Figure 3: Characterization of negative selection circuit and competition assay. a,** Plaque assay of an SP encoding AraC propagated on *E. coli* S2060 containing the $AP_{Prop}$ plasmid. IPTG was provided at increasing concentrations (0-100 µM) to induce expression of gene III. **b,** Expanded view of the competition assay from Fig. 2i. Experimental groups include individual infections and co-infections with WT, R2, and R5 phages, propagated in presence or absence of 10 mM L-arabinose. Cultures were supplemented with 25 µM IPTG at the time of phage infection and contained both the propagation ($AP_{Prop}$) and negative selection ($AP_{Neg}$) circuits. **c,** Sequence alignment of AraC transgenes from phages derived from the competition assay (see Fig. 2i). The L156I mutation, unique to the R5 variant, was used to identify R5-derived plaques.

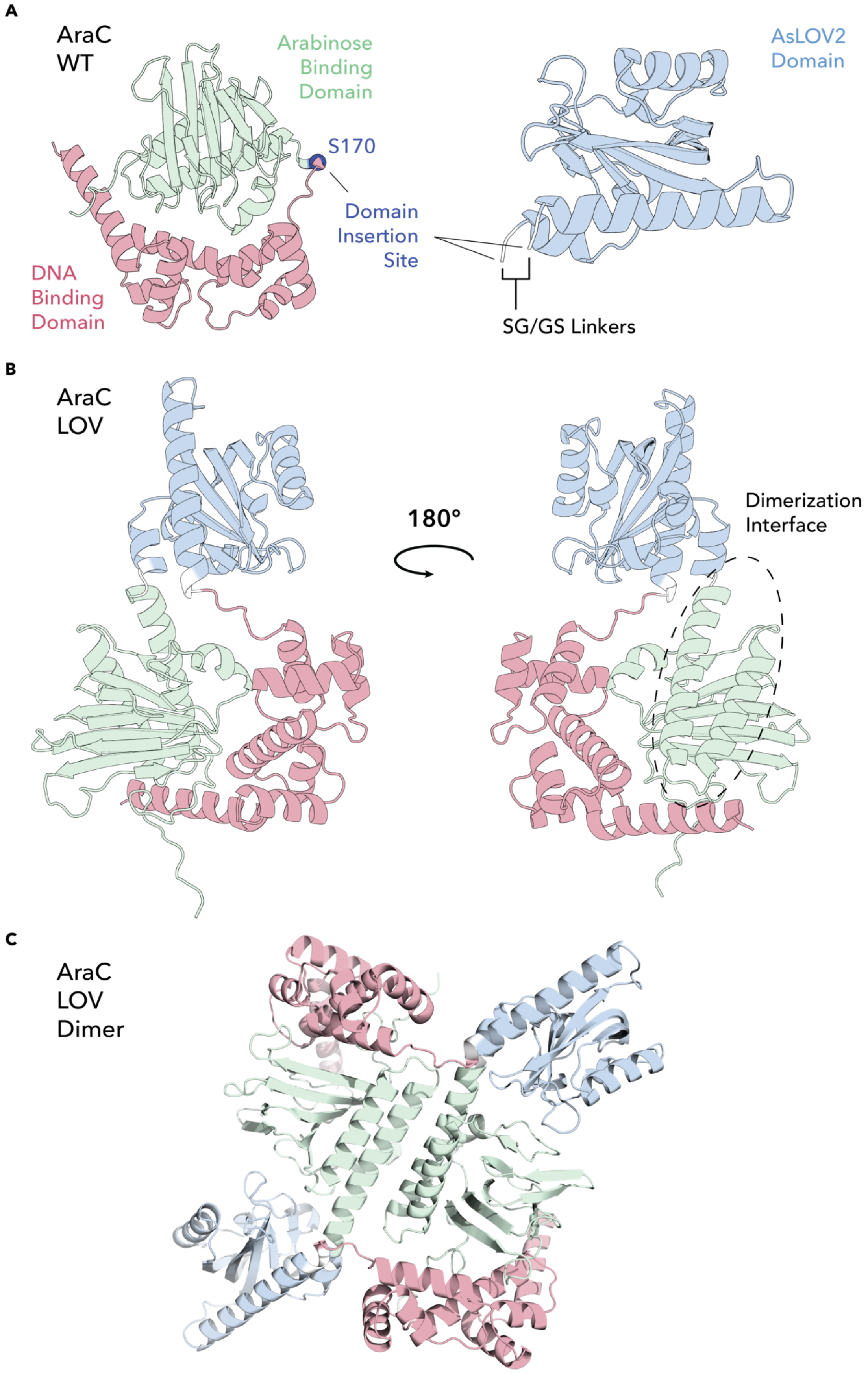
A
AraC WT
Arabinose Binding Domain
AsLOV2 Domain
S170
Domain Insertion Site
DNA Binding Domain
SG/GS Linkers
B
AraC LOV
180°
Dimerization Interface
C
AraC LOV Dimer

**Supplementary Figure 4. Structural context of the AraC–LOV domain insertion**. **a,** AlphaFold3 model of WT AraC. The arabinose-binding domain is shown in green and the DNA-binding domain in pink. The S170 domain-insertion site is marked with a blue sphere. The *As*LOV2 domain (blue) and the SG/GS linker segments (white) used in the AraC (WT)–LOV fusion are shown separately. **b,** AlphaFold3 model of WT–LOV, displayed in two orientations (180° rotation) using the same domain coloring as in panel A. The dimerization interface of AraC is circled with a dashed line. **c,** AlphaFold3-predicted WT-LOV dimer. A full assessment of all model confidence scores, including pLDDT, PAE, and interface-confidence metrics is provided in Supplementary Fig. 21.

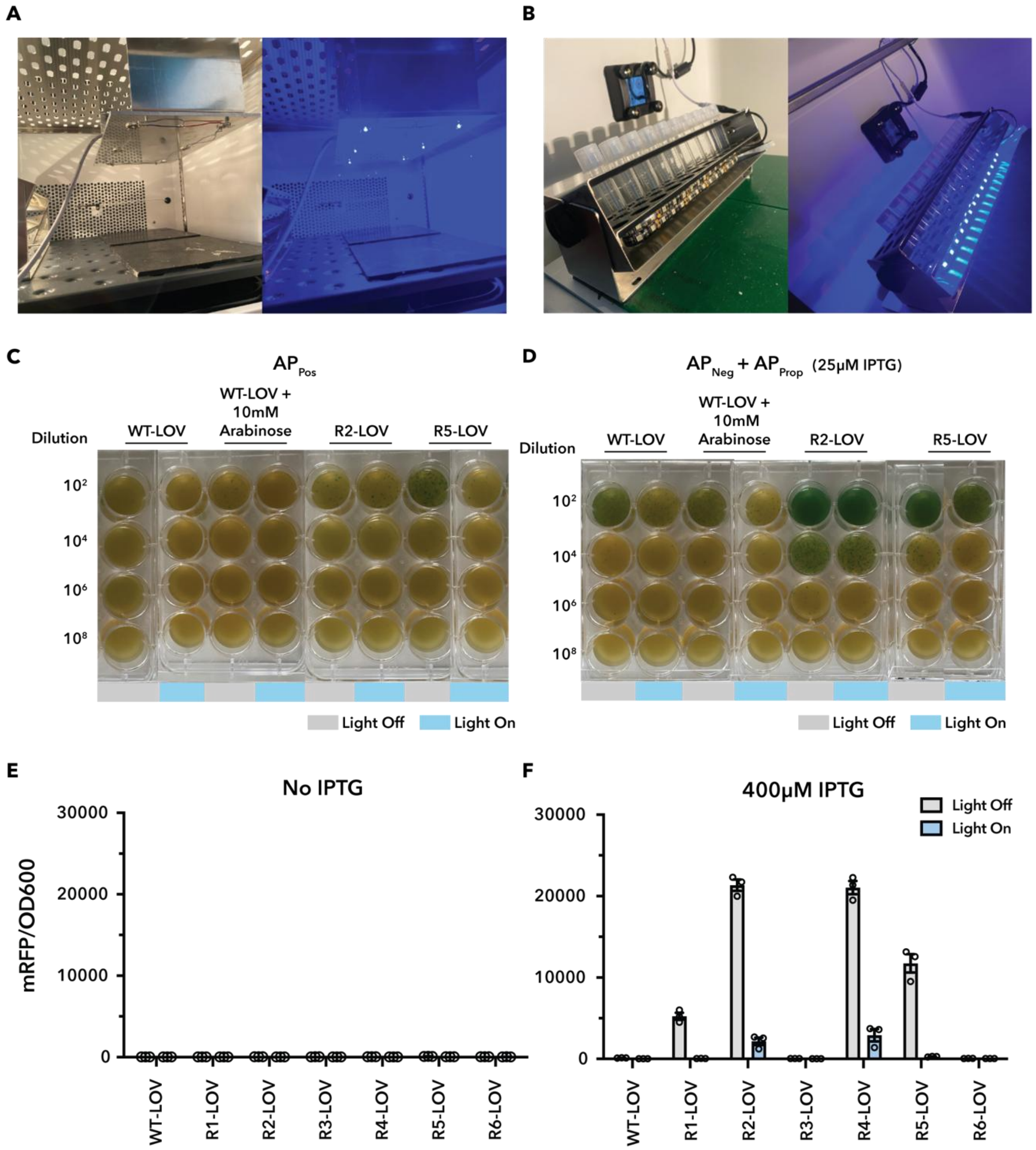


**Supplementary Figure 5: Functional assessment of AraC-LOV fusions based on evolved AraC variants. a,b** Overview of experimental blue light setups used for illumination of samples in **a,** multi-well plates or **b,** liquid cultures **c,** Plaque assay of phages carrying domain-inserted variants (WT-LOV, R2-LOV, R5-LOV) propagated under light or dark conditions in *E. coli* carrying the positive selection circuit (AP$_{Pos}$). A condition including WT-LOV with 10 mM L-arabinose is also shown. **d,** Experimental setup as in **c**, with the difference that cultures carry the negative selection circuit (APpos) and the propagation circuit (APprop), and were supplemented with 25 µM IPTG at the time of phage infection. **e,** RFP reporter assay of WT-LOV fusions based on WT AraC or the evolved R1-R6 variants expressed under sd2 RBS control, without IPTG induction. Cultures were incubated in the light or dark for 18 hours, followed by measurement of mRFP fluorescence and $OD_{600}$ in a plate reader. **f,** Experimental setup as in **e**, with the difference that cultures were induced with 400 μM IPTG. **e,f** Bars show mean ± s.e.m. of n = 3 biological replicates.

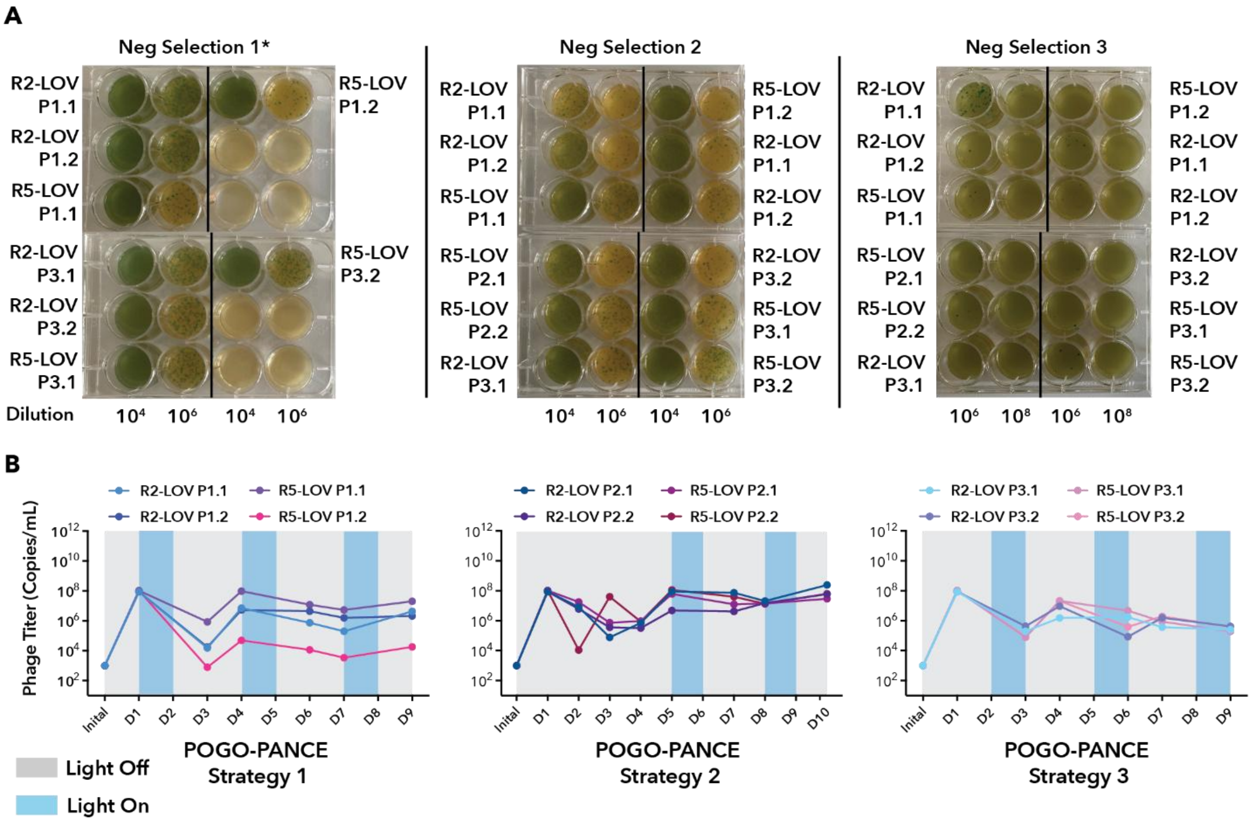


**Supplementary Figure 6: Tracking of POGO-PANCE evolution campaigns.**
**a,** Daily plaque assays from the negative selection step of each POGO-PANCE group. All rounds of negative selection are shown for R2-LOV and R5-LOV populations across the three evolutionary strategies. *P2.1 and 2.2 groups begin in Neg Selection group 2. **b,** Quantitative PCR analysis of phage titers during mutagenesis and positive selection phases for each POGO-PANCE pool. Titer trajectories under each strategy are shown separately.

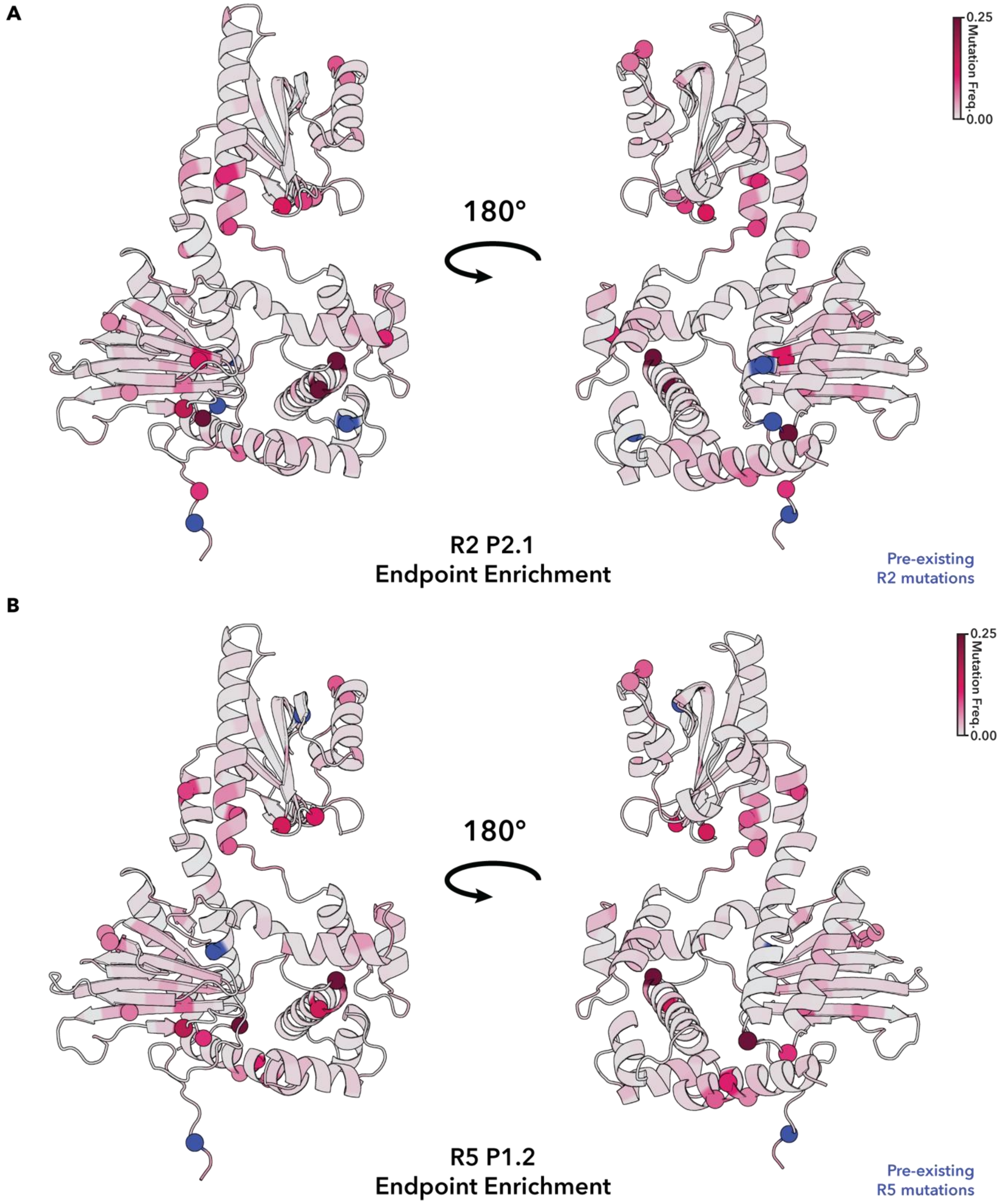


**Supplementary Figure 7: Endpoint mutation frequencies from long read sequencing are mapped onto Alphfold3-predicted structures of AraC-LOV for the indicated POGO-PANCE pools. a,** R2-LOV P2.1 pool. **b,** R5-LOV P1.2 pool.

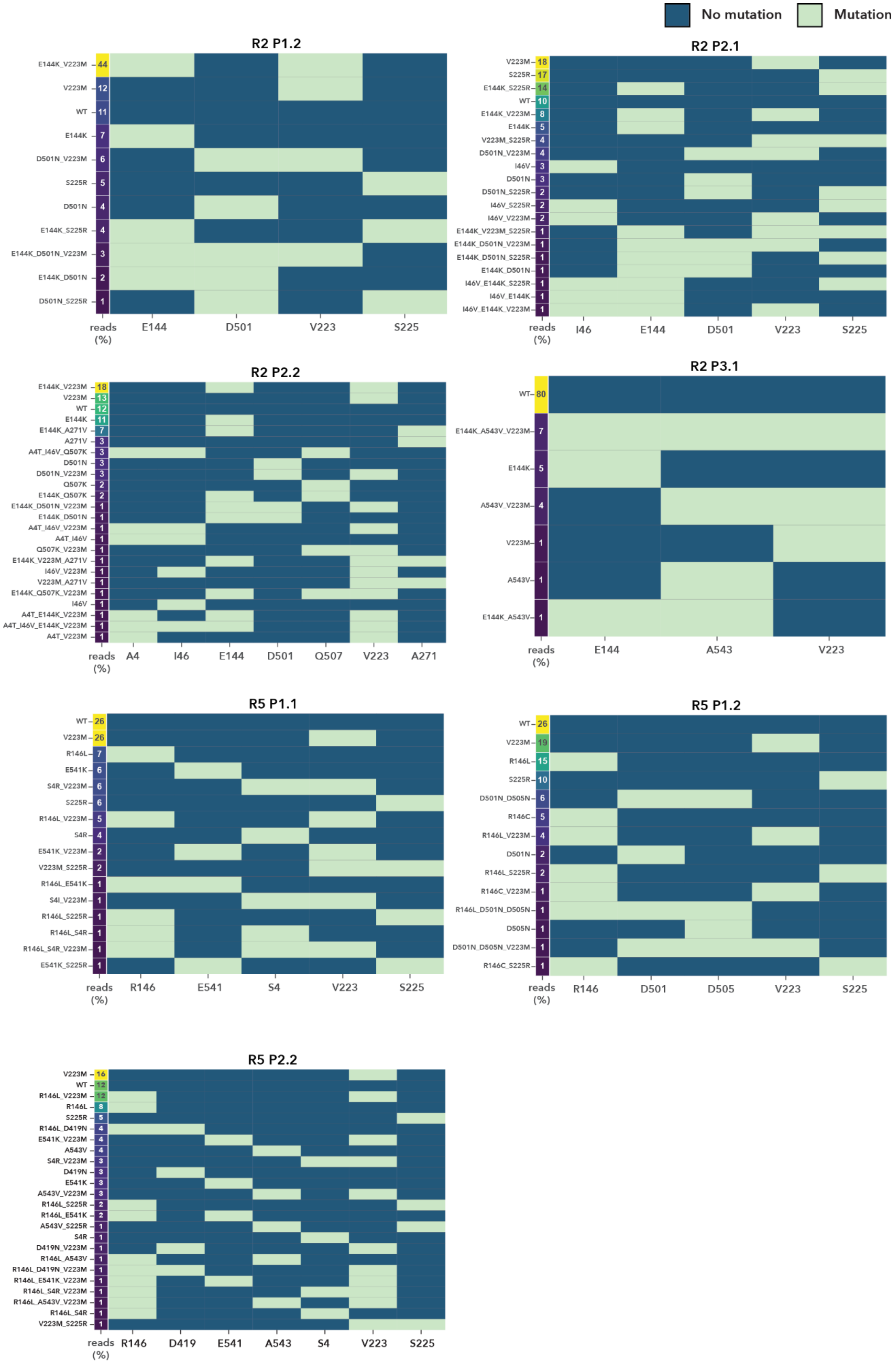
No mutation
Mutation
R2 P1.2
E144K_V223M 44
V223M 12
WT 11
E144K 7
D501N_V223M 6
S225R 5
D501N 4
E144K_S225R 4
E144K_D501N_V223M 3
E144K_D501N 2
D501N_S225R 1
reads (%)
E144
D501
V223
S225
R2 P2.1
V223M 18
S225R 17
E144K_S225R 14
WT 10
E144K_V223M 8
E144K 5
V223M_S225R 4
D501N_V223M 4
I46V 3
D501N 3
D501N_S225R 2
I46V_S225R 2
I46V_V223M 2
E144K_V223M_S225R 1
E144K_D501N_V223M 1
E144K_D501N_S225R 1
E144K_D501N 1
I46V_E144K_S225R 1
I46V_E144K 1
I46V_E144K_V223M 1
reads (%)
I46
E144
D501
V223
S225
R2 P2.2
E144K_V223M 18
V223M 13
WT 12
E144K 11
E144K_A271V 7
A271V 3
A4T_I46V_Q507K 3
D501N 3
D501N_V223M 3
Q507K 2
E144K_Q507K 2
E144K_D501N_V223M 1
E144K_D501N 1
A4T_I46V_V223M 1
A4T_I46V 1
Q507K_V223M 1
E144K_V223M_A271V 1
I46V_V223M 1
V223M_A271V 1
E144K_Q507K_V223M 1
I46V 1
A4T_E144K_V223M 1
A4T_I46V_E144K_V223M 1
A4T_V223M 1
reads (%)
A4
I46
E144
D501
Q507
V223
A271
R2 P3.1
WT 80
E144K_A543V_V223M 7
E144K 5
A543V_V223M 4
V223M 1
A543V 1
E144K_A543V 1
reads (%)
E144
A543
V223
R5 P1.1
WT 26
V223M 26
R146L 7
E541K 6
S4R_V223M 6
S225R 6
R146L_V223M 5
S4R 4
E541K_V223M 2
V223M_S225R 2
R146L_E541K 1
S4I_V223M 1
R146L_S225R 1
R146L_S4R 1
R146L_S4R_V223M 1
E541K_S225R 1
reads (%)
R146
E541
S4
V223
S225
R5 P1.2
WT 26
V223M 19
R146L 15
S225R 10
D501N_D505N 6
R146C 5
R146L_V223M 4
D501N 2
R146L_S225R 2
R146C_V223M 1
R146L_D501N_D505N 1
D505N 1
D501N_D505N_V223M 1
R146C_S225R 1
reads (%)
R146
D501
D505
V223
S225
R5 P2.2
V223M 16
WT 12
R146L_V223M 12
R146L 8
S225R 5
R146L_D419N 4
E541K_V223M 4
A543V 4
S4R_V223M 3
D419N 3
E541K 3
A543V_V223M 3
R146L_S225R 2
R146L_E541K 2
A543V_S225R 1
S4R 1
D419N_V223M 1
R146L_A543V 1
R146L_D419N_V223M 1
R146L_E541K_V223M 1
R146L_S4R_V223M 1
R146L_A543V_V223M 1
R146L_S4R 1
V223M_S225R 1
reads (%)
R146
D419
E541
A543
S4
V223
S225

**Supplementary Figure 8: Overview of enriched endpoint mutations in evolved phage populations.** Co-occurrence of mutations in endpoint phage pools was analyzed using long-read Nanopore sequencing. To mitigate the impact of sequencing noise, only positions with mutation enrichment exceeding 5% were included in the analysis, with the exception of position R38, which was excluded due to systematic errors observed across all pools. Pools lacking any positions above this threshold were omitted from the co-occurrence analysis. Within each qualifying pool, only variants present in more than 0.5% of total reads are displayed. Each heatmap depicts the frequency of these genotypes alongside their constituent mutations.

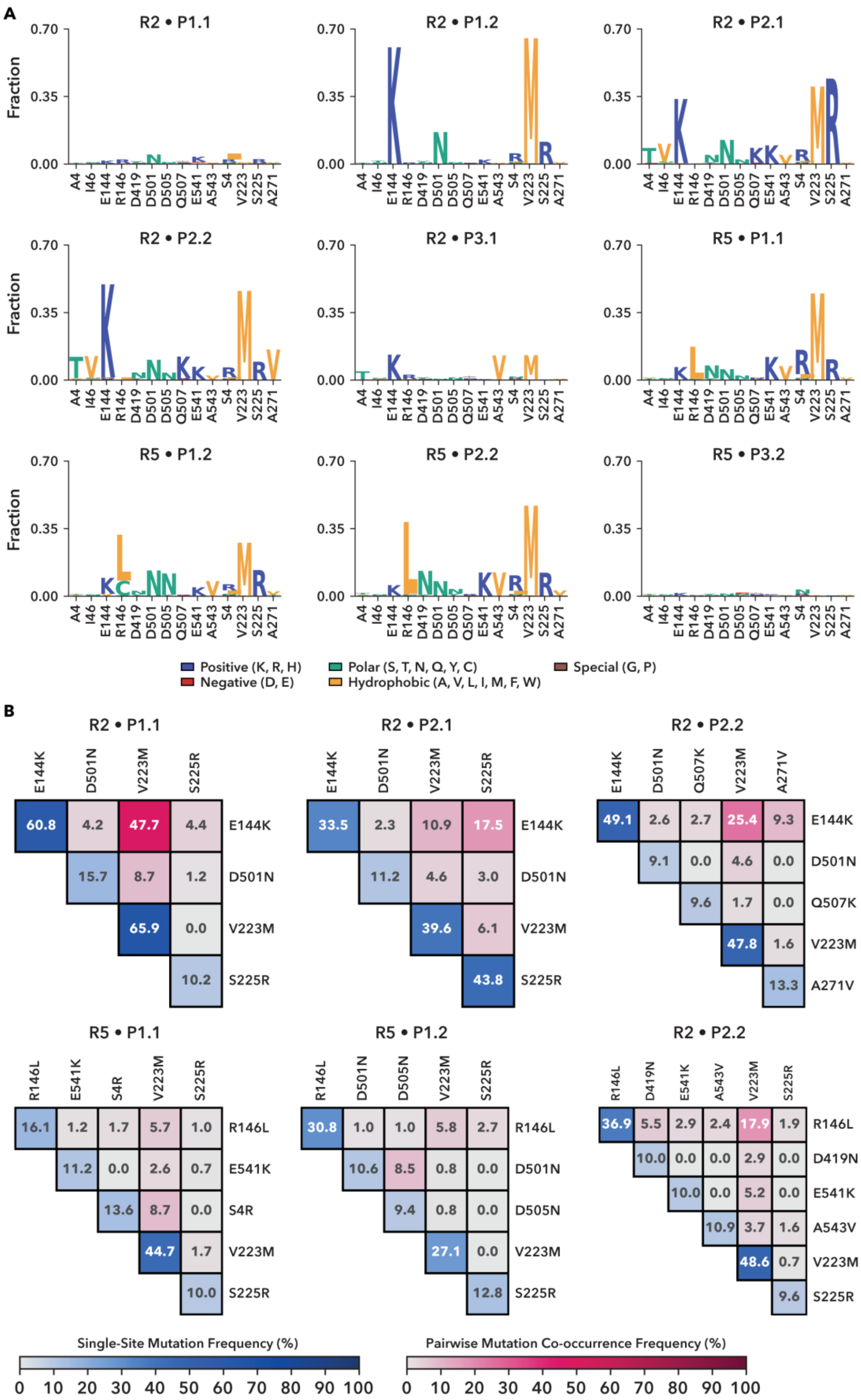
A
R2 • P1.1
R2 • P1.2
R2 • P2.1
R2 • P2.2
R2 • P3.1
R5 • P1.1
R5 • P1.2
R5 • P2.2
R5 • P3.2
Fraction
Positive (K, R, H)
Negative (D, E)
Polar (S, T, N, Q, Y, C)
Hydrophobic (A, V, L, I, M, F, W)
Special (G, P)
B
Single-Site Mutation Frequency (%)
Pairwise Mutation Co-occurrence Frequency (%)

**Supplementary Figure 9: Analysis of residue level enrichment in the final day POGO-PANCE pools. a,** Sequence logo plots showing the fraction of the most highly enriched amino acid at each mutated position (enrichments above 1% across the final-day pools were considered in the analysis). Colors correspond to amino acid classes as indicated in the legend (positive, negative, polar, hydrophobic, and special). **b,** Pairwise mutation co-occurrence matrices are shown for each pool. Blue squares indicate the single-site mutation frequency of each residue in the pool, and burgundy squares indicate the pairwise co-occurrence frequency between residues within individual reads. Only residues enriched in at least 9.5% of reads are included.

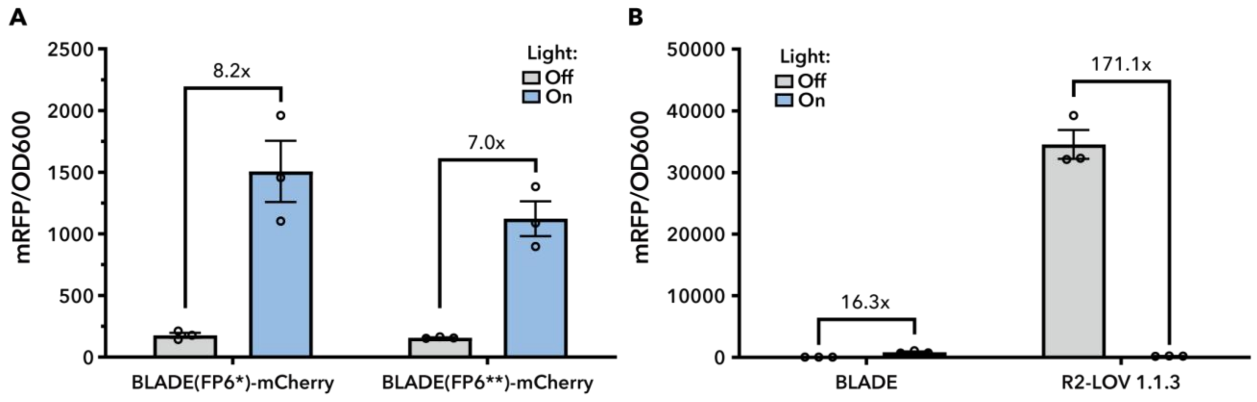


**Supplementary Figure 10: Comparison of BLADE system with R2-LOV 1.1.3**
**a,** RFP reporter assay BLADE plasmids sourced from the top candidates from Romano et. al. Cultures were incubated in the light or dark for 18 hours, followed by measurement of mRFP fluorescence and $OD_{600}$ in a plate reader. **b**, BLADE and R2-LOV 1.1.3 expressed under sd2 RBS control. Experimental setup as in **A**, with the difference that cultures were induced with 400 μM IPTG. **a,b,** Bars show mean ± s.e.m. of n = 3 biological replicates.

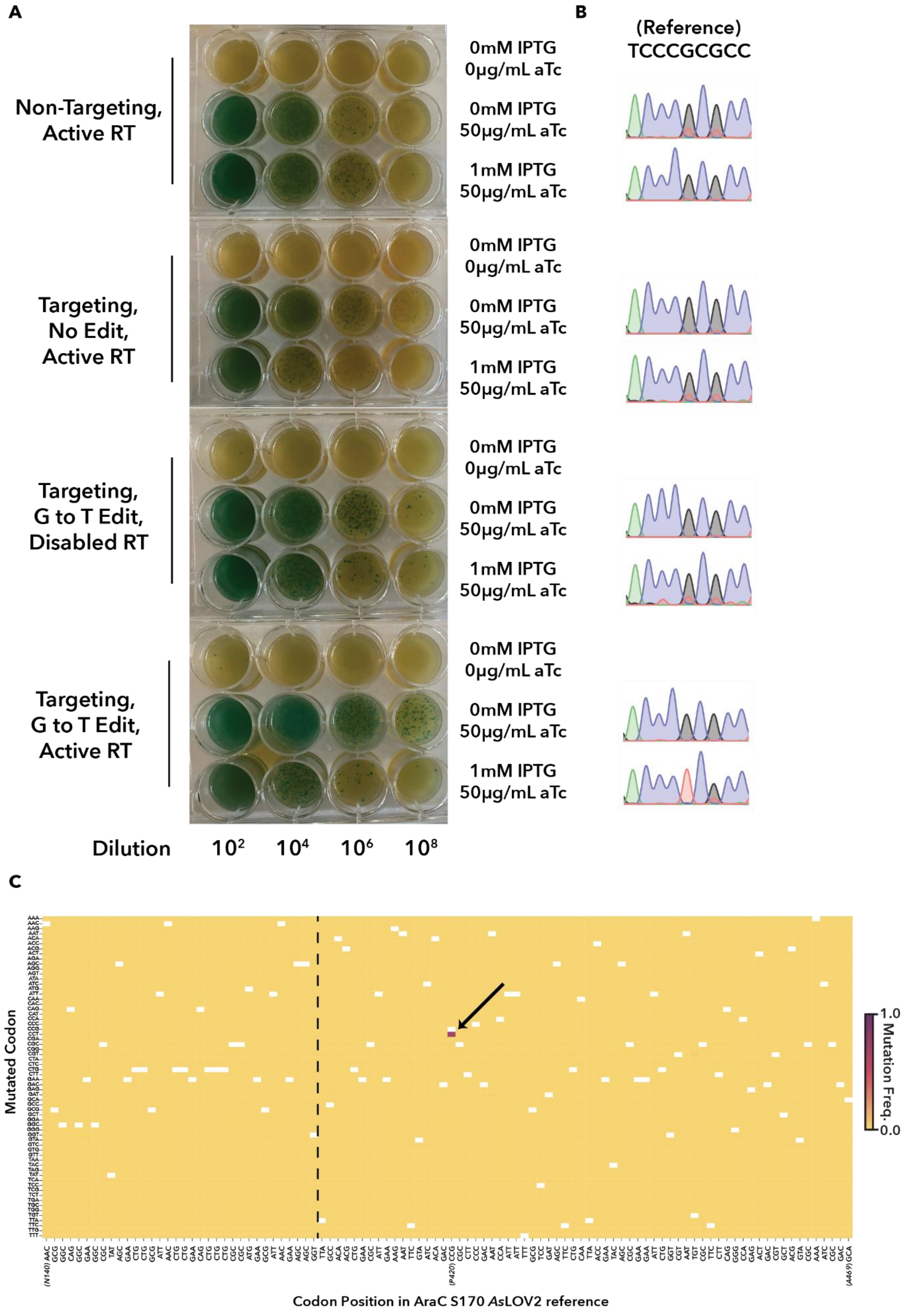
A
Non-Targeting,
Active RT
Targeting,
No Edit,
Active RT
Targeting,
G to T Edit,
Disabled RT
Targeting,
G to T Edit,
Active RT
0mM IPTG
0µg/mL aTc
0mM IPTG
50µg/mL aTc
1mM IPTG
50µg/mL aTc
Dilution
10²
10⁴
10⁶
10⁸
B
(Reference)
TCCCGCGCC
C
Mutated Codon
Codon Position in AraC S170 AsLOV2 reference
(N140)
(P420)
(A469)
1.0
0.0
Mutation Freq.

**Supplementary Figure 11: Validation of retron-based editing using a single substitution retron. a,** Plaque assay comparing editing outcomes from different Recombitron configurations. Conditions include: a non-targeting retron with active reverse transcriptase (RT); a targeting retron with no edit (active RT); a targeting retron encoding a G-to-T substitution, but carrying a catalytically inactive RT; and a targeting retron with the G-to-T substitution and active RT. Plaque output was assessed after infection and full induction with aTc (for gIII production) and IPTG (for Recombitron expression). **b,** Sanger sequencing chromatograms of phage pools from each aTc-induced condition in **a**. Efficient editing was observed only in the presence of an active RT and an edit-encoding retron, resulting in a fully converted allele with no detectable wild-type sequence. **c,** Nanopore sequencing of phage populations edited with a retron encoding a silent G-to-T substitution (CCG→CCT) at LOV residue P420 of AraC-LOV, as in panels **a** and **b**. Using a $10^{10}$ pfu/mL input, 71% editing was achieved at the target site (arrow), while no editing is observed elsewhere. Codons are plotted by reference position (x-axis) and variant identity (y-axis), with the first, final and edited site marked and the start of LOV denoted with a dashed line.

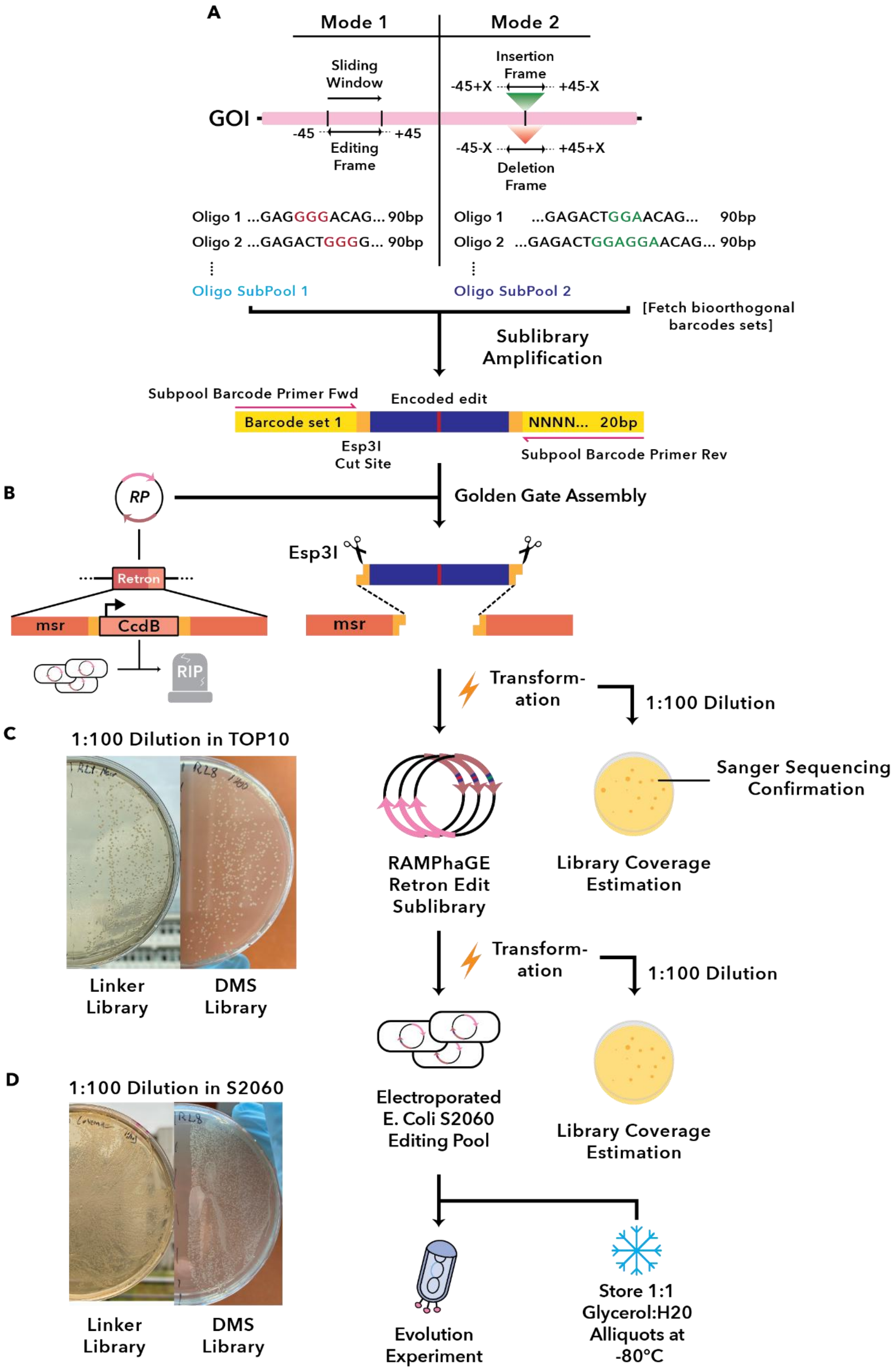

A
Mode 1
Mode 2
Sliding Window
Insertion Frame
-45+X
+45-X
GOI
-45
+45
Editing Frame
-45-X
+45+X
Deletion Frame
Oligo 1 ...GAGGGGACAG... 90bp
Oligo 2 ...GAGACTGGGG... 90bp
Oligo 1 ...GAGACTGGAACAG... 90bp
Oligo 2 ...GAGACTGGAGGAACAG... 90bp
Oligo SubPool 1
Oligo SubPool 2
[Fetch bioorthogonal barcodes sets]
Sublibrary Amplification
Subpool Barcode Primer Fwd
Encoded edit
Barcode set 1
NNNN... 20bp
Esp3I Cut Site
Subpool Barcode Primer Rev
B
RP
Golden Gate Assembly
Retron
msr
CcdB
RIP
Esp3I
msr
Transform-ation
1:100 Dilution
C
1:100 Dilution in TOP10
Sanger Sequencing Confirmation
RAMPhaGE Retron Edit Sublibrary
Library Coverage Estimation
Linker Library
DMS Library
Transform-ation
1:100 Dilution
D
1:100 Dilution in S2060
Electroporated E. Coli S2060 Editing Pool
Library Coverage Estimation
Linker Library
DMS Library
Evolution Experiment
Store 1:1 Glycerol:H20 Alliquots at -80°C

**Supplementary Figure 12: Construction of barcoded retron recombineering libraries for phage genome editing. a,** Schematic overview of retron oligonucleotide design and barcoding. Retron donor oligos were designed either as overlapping 90-bp sliding windows across the gene of interest (DMS library) or as fixed-position sequences encoding predefined substitutions, insertions, or deletions (Linker library). Each oligo included a maximum of 45 bp homology arms flanking the central edit, autogenerated in both forward and reverse directions. Oligo subpools were PCR-amplified using primers bearing unique bioorthogonal barcodes and flanked by Esp3I restriction sites. **B,** Cloning of retron oligo subpools into the retron expression plasmid. Amplified subpools were inserted into a retron plasmid backbone via Golden Gate assembly, replacing a toxic *ccdB* cassette to enable positive selection in *E. coli* TOP10. **c,** Assembly validation and transformation. Retron library plasmids were transformed into TOP10 cells, cultured overnight, and miniprepped. A fraction of each transformation was plated to estimate library coverage, and selected colonies were verified by Sanger sequencing. **d,** Preparation of phage editing host cell libraries. Validated retron plasmid libraries were transformed into *E. coli* S2060 cells. A fraction was plated to quantify editing library coverage. Cell pools were either used directly in phage editing experiments or stored at −80 °C in 1:1 LB and 50% glycerol.

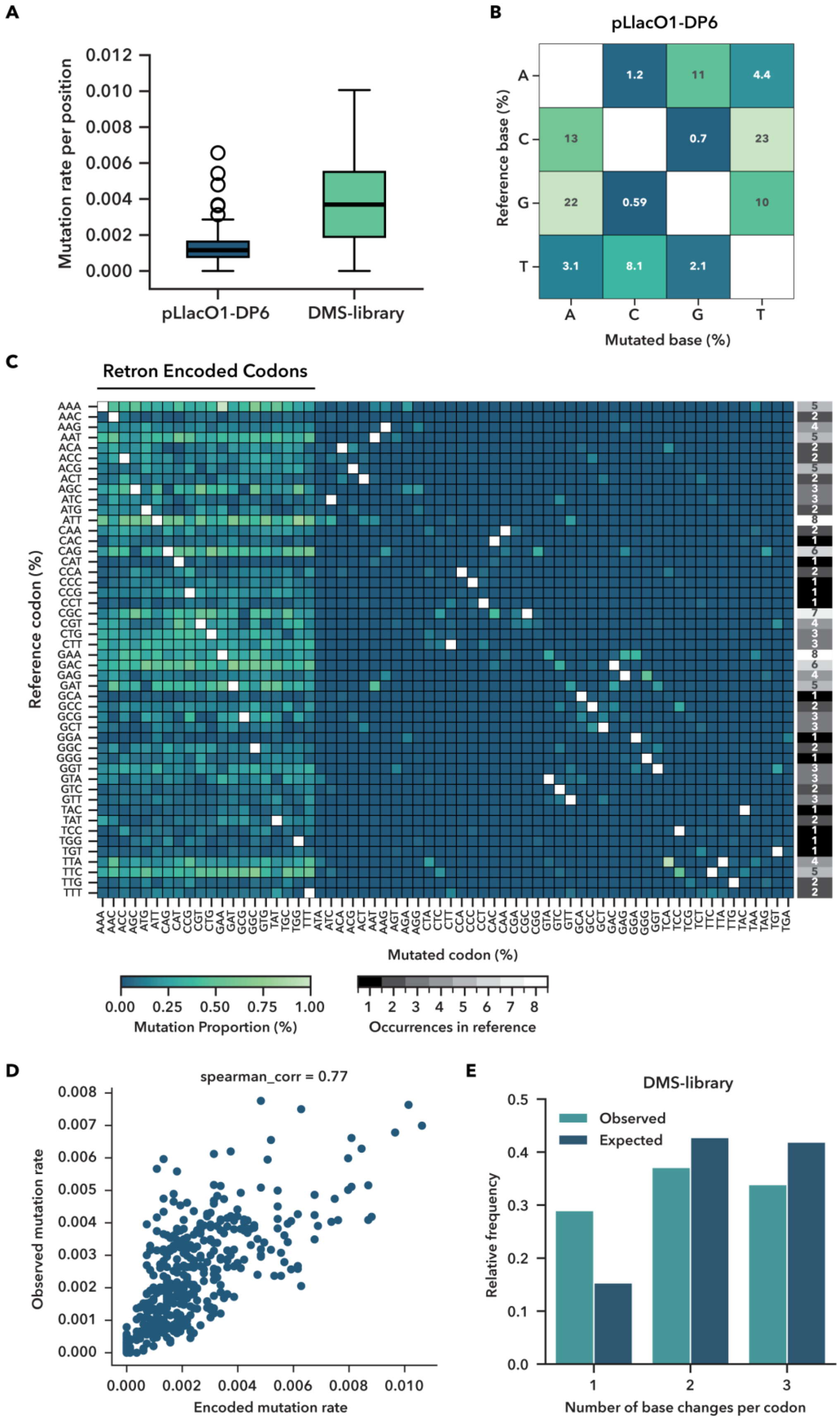
A
Mutation rate per position
0.012
0.010
0.008
0.006
0.004
0.002
0.000
pLlacO1-DP6
DMS-library
B
pLlacO1-DP6
Reference base (%)
Mutated base (%)
A
C
G
T
1.2
11
4.4
13
0.7
23
22
0.59
10
3.1
8.1
2.1
C
Retron Encoded Codons
Reference codon (%)
Mutated codon (%)
0.00 0.25 0.50 0.75 1.00
Mutation Proportion (%)
1 2 3 4 5 6 7 8
Occurrences in reference
D
spearman_corr = 0.77
Observed mutation rate
Encoded mutation rate
0.000 0.002 0.004 0.006 0.008 0.010
E
DMS-library
Observed
Expected
Relative frequency
Number of base changes per codon
1
2
3

**Supplementary Figure 13: Comparative analysis of mutational spectra and coverage between DP6 mutagenesis and retron-encoded DMS libraries. a-e,** AraC-LOV encoding phage was propagated overnight either on pLlacO1-DP6 or the retron DMS library containing host cells, followed by PCR of the Acr-LOV fragment from phage pools and subsequent NGS. **a,** Total amino acid mutation rates per site were calculated as the sum of all normalized mutation frequencies for the LOV domain. The DMS library exhibited ~threefold higher median per-position mutation rates compared to DP6, reflecting increased editing density. **b,** Nucleotide-level mutational spectrum of plLacO1-DP6 shows strong biases well-known for the parent mutagenesis construct MP6. **c,** Codon substitution spectrum in the DMS-edited phage population. Heatmap shows the relative frequency (%) of all observed codon substitutions in the LOV region as normalized per position. Columns indicate codons encoded in the DMS library ("Retron Encoded Codons") or absent, while rows represent all possible codon identities sorted alphabetically. The greyscale heatmap on the right show the number of times each codon occurs in the WT LOV reference sequence. **d,** Correlation of positional mutation frequencies between the DMS plasmid library and the phage populations obtained upon propagation on respective hosts. Relative frequencies of plasmid-encoded versus actual phage edited nucleotide substitutions were compared per position. A strong correlation was observed (Spearman $\rho = 0.77$), indicating that the positional distribution of observed mutations is largely determined by the composition of the retron-encoding plasmid library. **e,** Codon edits recovered from DMS-edited phages were categorized according to the number of nucleotide substitutions required to generate each amino acid change (one-, two-, or three-nucleotide substitutions), and the proportion of each class was calculated across all edited LOV codons. Expected proportions (blue) were obtained by applying the same classification to every codon substitution encoded in the DMS library, assuming equal likelihood of sampling each variant. These expected values were compared to the observed proportions (green).

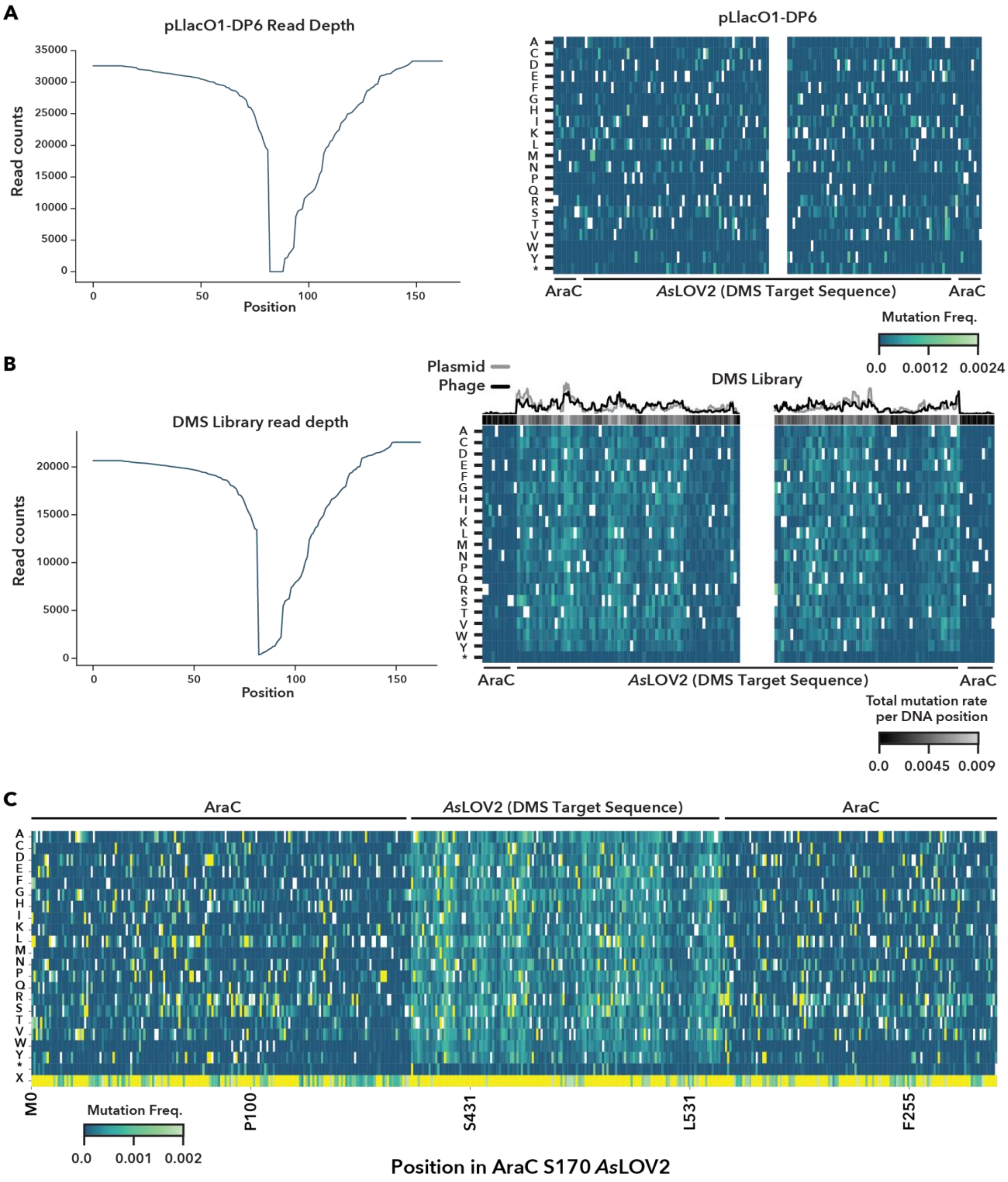


**Supplementary Figure 14: Comparison of mutation rates, positional biases, and sequencing coverage between DP6 mutagenesis and the retron-encoded DMS library. a-c** AraC-LOV encoding phage was propagated overnight either on pLlacO1-DP6 or the retron DMS library containing host cells, followed by PCR of the AraC WT-LOV fragment from phage pools and subsequent NGS. **a,** Illumina-based analysis of mutagenesis by plLacO1-DP6. Left: Read coverage across the reference sequence (x-axis, DNA level) after filtering for quality, indels, and low coverage; y-axis shows read counts. Coverage drops toward the middle of the sequence, with an average coverage of 75.26% across the region of interest. Right: Mutation frequencies are shown per amino acid position across WT-LOV. Each heatmap cell reflects the normalized frequency of non-reference amino acids at that position. White cells indicate reference residues or positions with read depth <2,000. **b,** llumina-based assessment of DMS library editing. Left: Read coverage after quality filtering, with an average coverage of 72.84% across the region. Right: Bottom heatmap (blue) shows observed amino acid mutation frequencies per position in the phage population. Middle heatmap (gray) displays total mutation frequency per DNA position. White cells indicate reference residues or positions with read depth <2,000. Line plots on top compare the relative frequency of plasmid encoded mutations (gray) versus mutations observed in WT-LOV phage PCR amplicons (black) per DNA residue (see Methods). Shaded sections regions intersecting the two lines highlight areas of positional bias. **c,** Nanopore-based profiling of DMS-edited phage populations. Mutation frequencies per amino acid position are plotted across the WT-LOV sequence. White cells indicate reference residues. Cells with mutation frequencies exceeding 0.002 are shaded in yellow; Due to Nanopore sequencing noise, the frequency scale was lowered to enhance signal detection. X refers to corrected positions with indels that would lead to a frameshift.

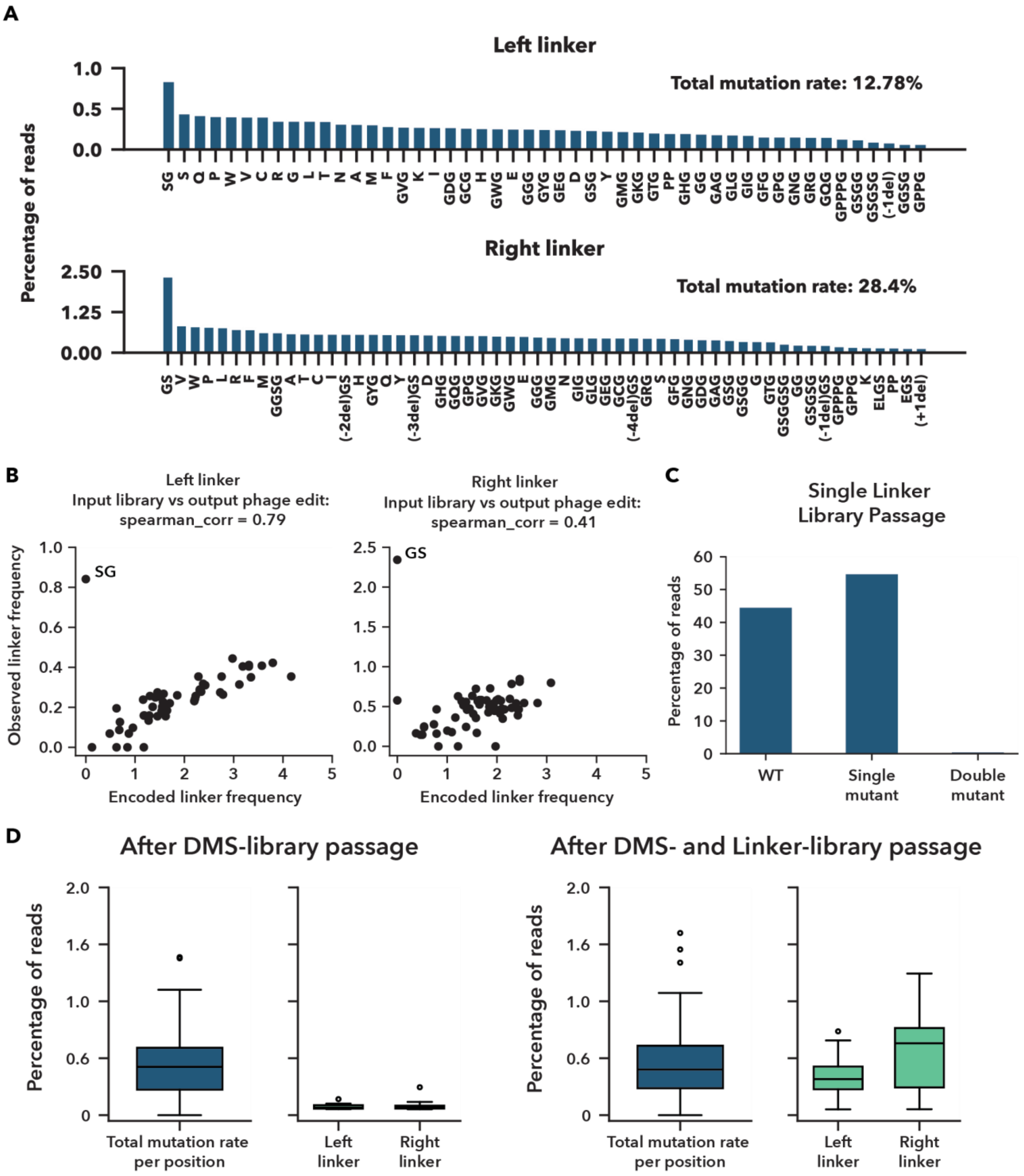


**Supplementary Figure 15: Mutational outcomes from retron-mediated editing of left and right linker regions. a,** Illumina-based analysis of linker variants after mutagenesis with the retron-encoded linker library. Variants include both on-target and functional off-target edits (e.g., silent mutations or indels that preserve reading frame). GS and SG linkers are included as WT on the amino acid level but carry synonymous mutations at the DNA level. Only variants representing >0.05% of reads are shown. **b,** Correlation between encoded and observed linker variants. Scatter plots compare the relative abundance of encoded linker variants in the retron-encoding plasmid input library to their observed frequency in edited phage populations. Left and right linker data are plotted separately. Spearman correlation values reflect the extent to which the observed mutational spectrum mirrors the library design. **c,** Distribution of WT and edited linker combinations after one passage of phage editing. Nanopore sequencing was used to categorize each phage as WT, single linker mutant, or double linker mutant based on the amino acid sequence. Variants occurring in <0.01% of reads were excluded to reduce noise. WT linkers include all sequences matching the reference amino acid sequence; silent DNA mutations were not considered. **d,** Combined effects of DMS and linker library editing. Phages were first propagated on host cells carrying the DMS retron library (left), followed by propagation on host cells containing the retron linker library (right). Total mutation frequency per position was calculated across the LOV insert. "IDEAAK" linker residues at the end of LOV were excluded from this DMS quantification, as they are encoded in the linker library. "Left Linker" and "Right Linker" correspond to the frequencies of individual linker variants before and after the subsequent application of the linker library. Data show the percentage of total reads containing each linker variant, including off-target edits and edits at overlapping positions between libraries. Variants occurring in <0.05% of reads were excluded. Only observed mutations are plotted; variants that were encoded but not detected are not shown.

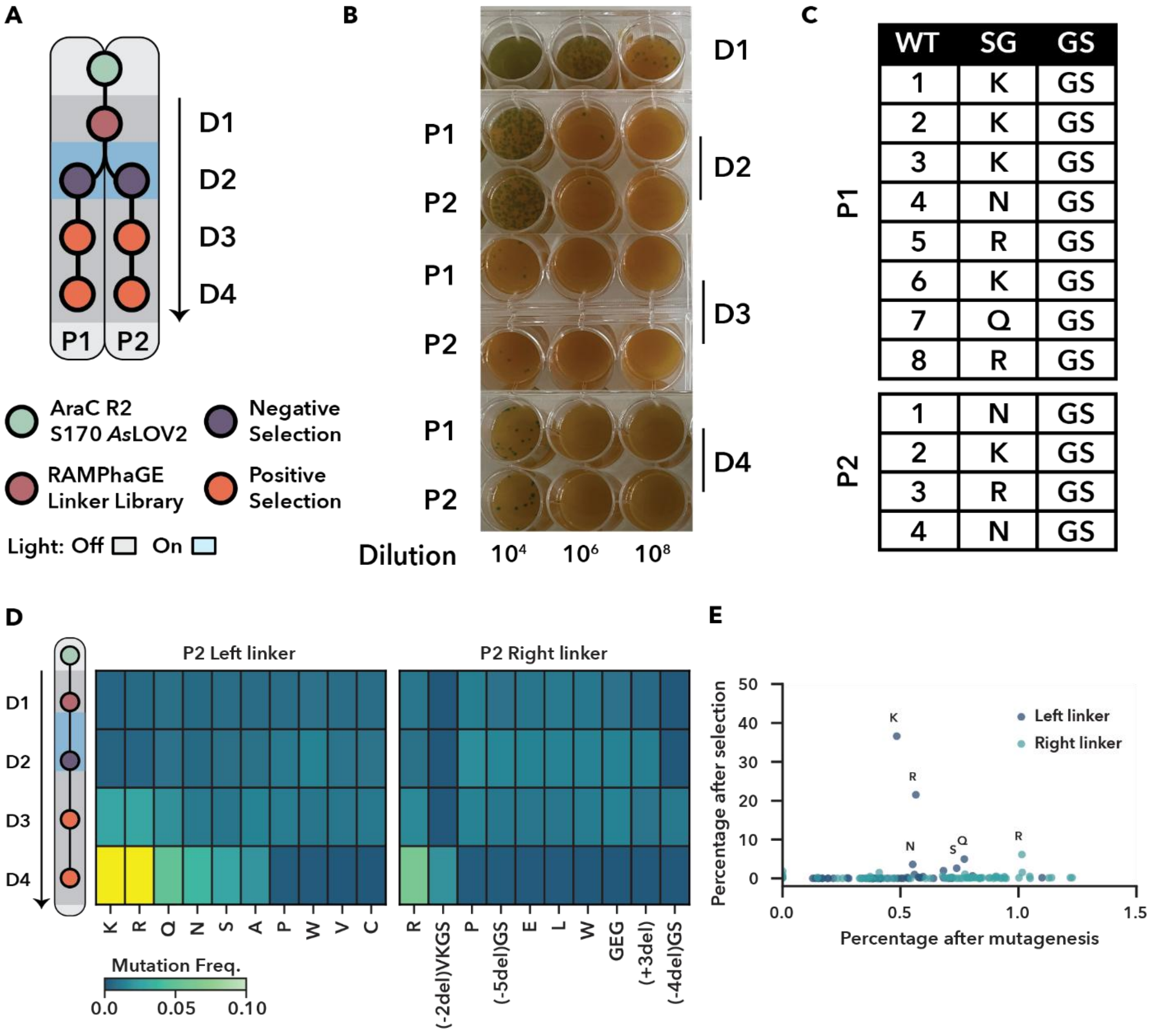


**Supplementary Figure 16: Expanded analysis of linker library evolution via POGO-PANCE. a,** Expanded schematic of the linker library POGO-PANCE evolution strategy from Fig 5g. Two parallel evolution pools (P1 and P2) were initiated from a shared RAMPhaGE-edited linker library based on the R2-LOV input phage. Each pool underwent iterative cycles of mutagenesis, negative selection, and positive selection under light and dark conditions, as indicated (see Methods for details). **b**, Plaque assays tracking phage propagation during each selection day across both P1 and P2. Dilution series are shown for all four days of mutagenesis and selection. Approximate log-scale titers for both evolution groups followed a similar trajectory of $10^{10}$, $10^8$, $10^6$, and $10^7$ pfu/mL across the four days. **c,** Genotypic analysis of individual plaques sampled from each pool at the end of the evolution. Left and right linker sequences are annotated, with WT and SG variants highlighted. Enriched linker residues (e.g., K, Q, N, R) are indicated. **d,** Temporal enrichment of linker variants in pool P2 during POGO-PANCE evolution. Enrichment was calculated as the mutation frequency per linker residue across four consecutive days, with residues exceeding 0.10 mutation frequency shaded in yellow. The top 10 linkers with the highest variance across all POGO-PANCE steps are shown for each site. **e,** Scatter plot comparing the percentage of each linker variant after one passage of

RAMPhaGE mutagenesis (x-axis) to the final enrichment after POGO-PANCE selection (y-axis) in pool P2. Select linker residues are labeled.

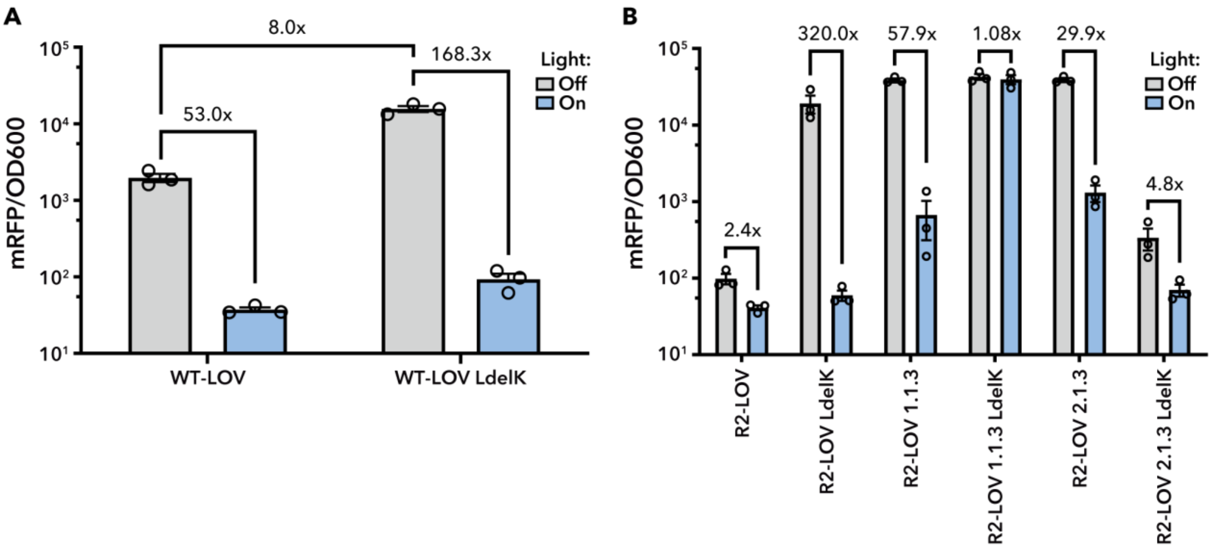


**Supplementary Figure 17: Transplantation of LdelK linker into different AraC–LOV variants. a,** RFP reporter assay comparing WT-LOV and WT-LOV LdelK. Cultures were induced with 10 mM L-arabinose and 400 µM IPTG, incubated in the light or dark for 18 h, and mRFP fluorescence and OD600 were measured in a plate reader. **b,** RFP reporter assay of R2-LOV variants expressed under sd2 RBS control. Experimental conditions were as in panel A, except that no L-arabinose was added. **a,b,** Bars show mean ± s.e.m. of n = 3 biological replicates.

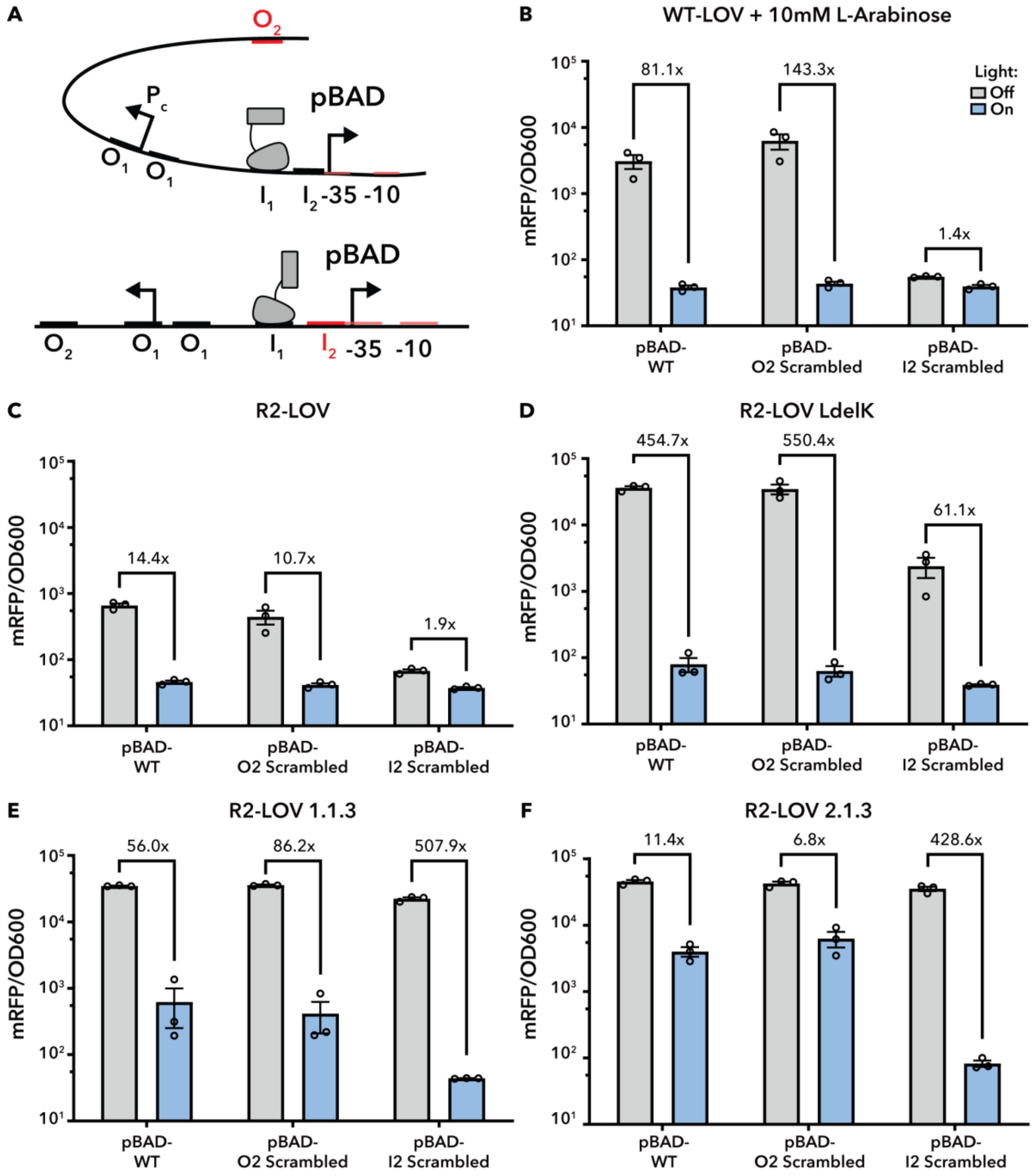


**Supplementary Figure 18: Effects of pBAD promoter modification on the performance of different AraC–LOV variants. a,** Schematics of the pBAD promoter showing either a scrambled $O_2$ half-site or $I_2$ half-site (the −35 element within $I_2$ preserved). **b-f,** RFP reporter assays for multiple AraC–LOV variants (indicated) using the WT pBAD promoter, an $O_2$-scrambled promoter, or an $I_2$-scrambled promoter. Cultures were induced with 400 µM IPTG, incubated in the light or dark for 18 h (and mRFP fluorescence and $OD_{600}$ were measured in a plate reader (bars show mean ± s.e.m.; n = 3 biological replicates). The light/dark legend shown in panel b also applies to panels c-f. **b,** WT-LOV induced with 10 mM L-arabinose. **c,** R2-LOV. **d,** R2-LOV LdelK. **e,** R2-LOV 1.1.3. **f,** R2-LOV 2.1.3.

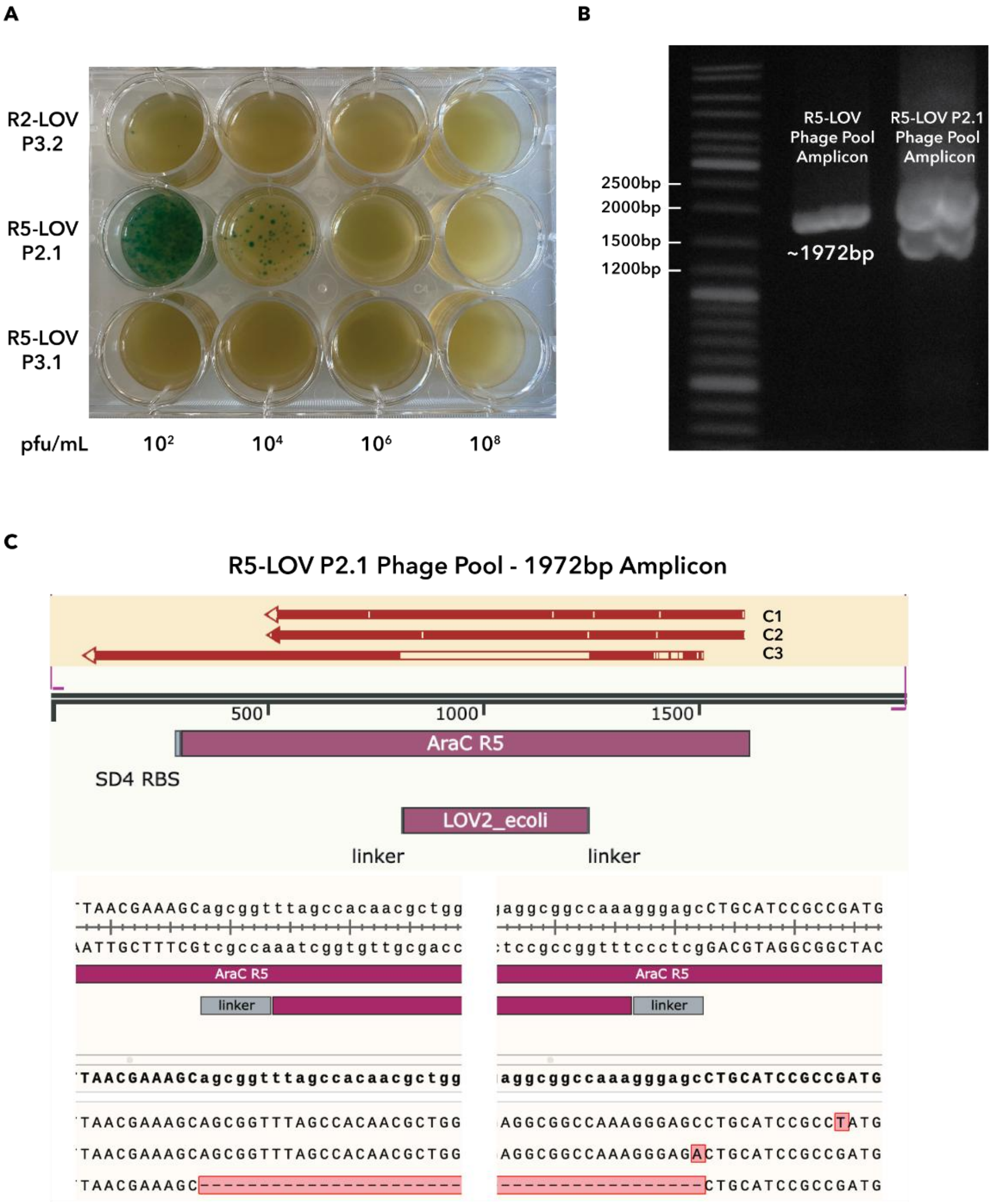


**Supplementary Figure 19: Failure modes observed in eliminated POGO-PANCE pools.**
**a,** Plaque assays of endpoint phage pools from R2-LOV P3.2, R5-LOV P2.1, and R5-LOV P3.1, showing minimal propagation for P3 pools. **b,** Agarose gel analysis of R5-LOV phage and the R5-LOV P2.1 endpoint pool amplicons. The expected size of the R5-LOV amplicon (1972 bp) is indicated. **c,** Alignment of sequencing reads from three plaques derived from the R5-LOV P2.1 endpoint pool to the R5-LOV reference sequence. Plaques C1 and C2 contain coding point mutations, whereas plaque C3 lacks the LOV domain entirely.

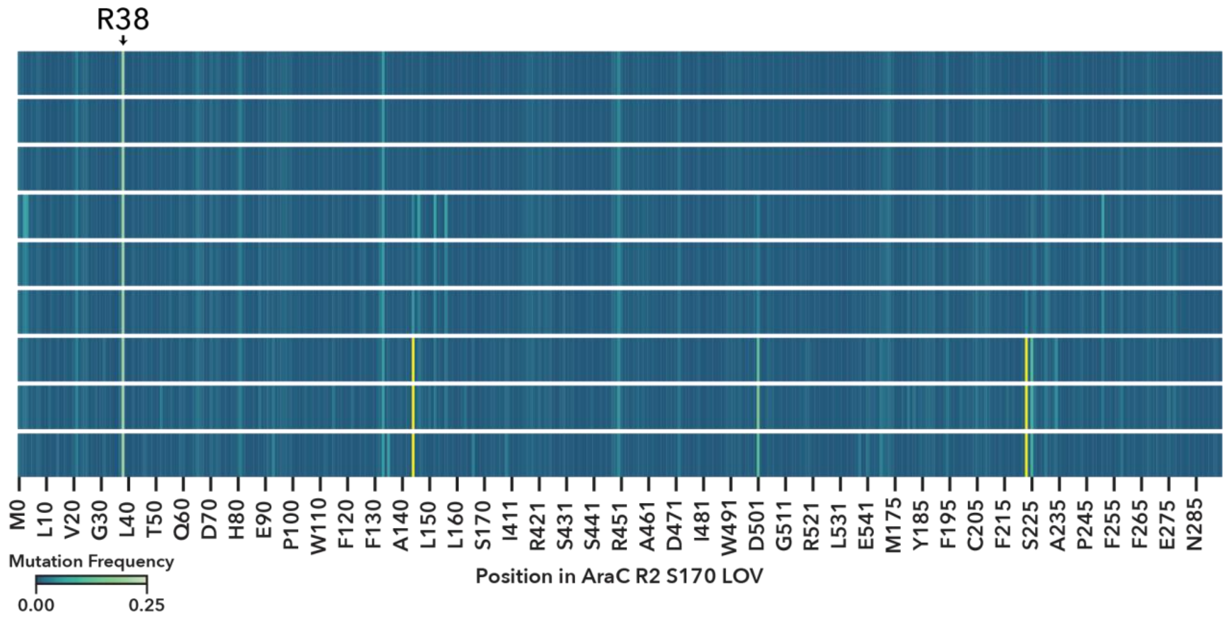


**Supplementary Figure 20: Uncorrected temporal mutation enrichment across the AraC-LOV R2 P1.2 sequence during POGO-PANCE evolution.** Nanopore-based tracking of mutation frequencies across the R2-LOV sequence over three complete cycles of mutagenesis, negative selection, and positive selection. Data correspond to the temporal enrichment analysis presented in Fig. 3c. Mutation frequencies are shown for each cycle by amino acid position, with residues exceeding 0.25 mutation frequency shaded in yellow. To reduce noise and correct for systematic sequencing artifacts, such as R38 (indicated), background mutation rates observed during the first mutagenesis cycle, assumed to reflect random mutagenesis plus persistent technical noise, were subtracted from all data sets in Fig. 3c.

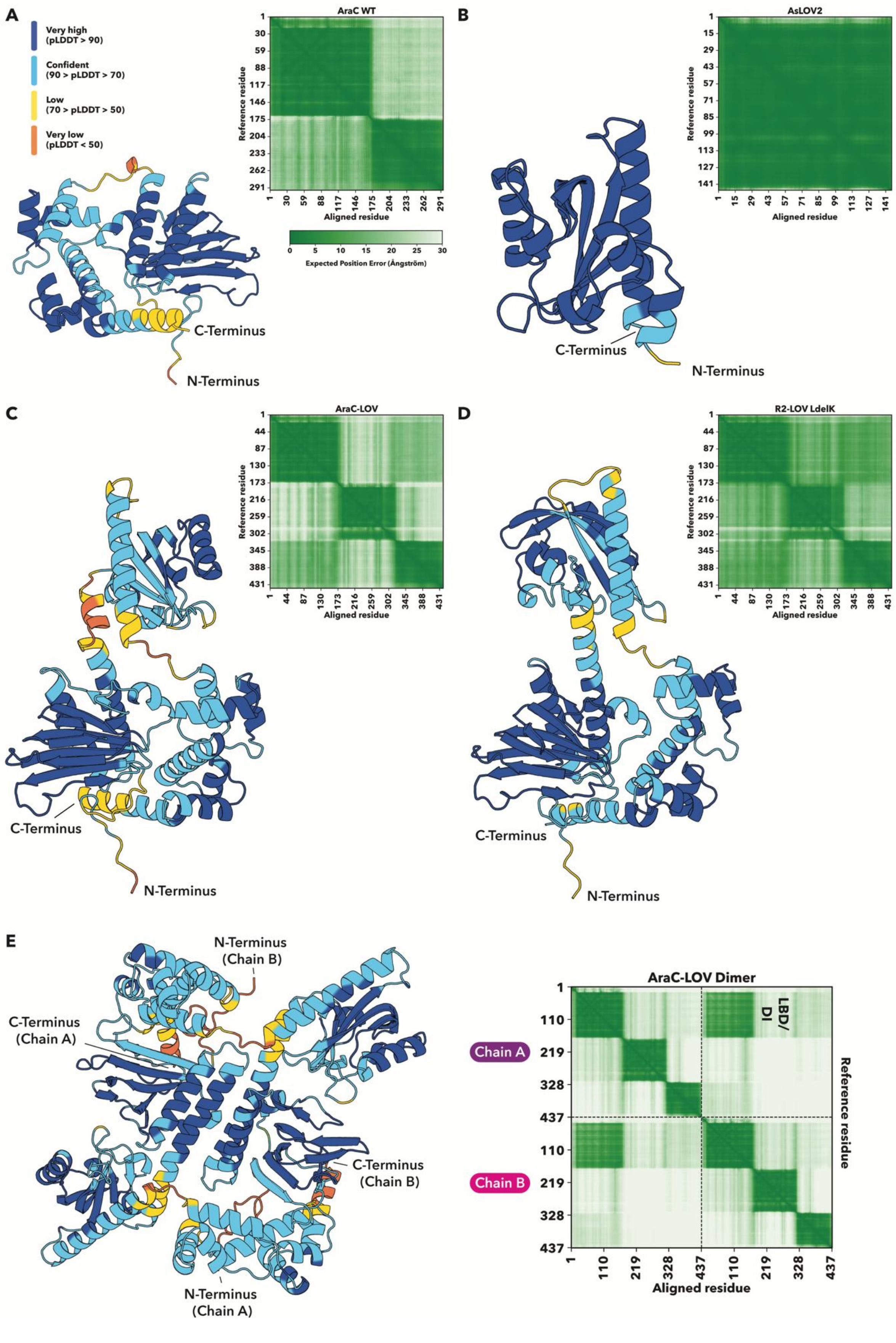
A
Very high (pLDDT > 90)
Confident (90 > pLDDT > 70)
Low (70 > pLDDT > 50)
Very low (pLDDT < 50)
AraC WT
Reference residue
Aligned residue
Expected Position Error (Ångström)
C-Terminus
N-Terminus
B
AsLOV2
C
AraC-LOV
D
R2-LOV LdelK
E
N-Terminus (Chain B)
C-Terminus (Chain A)
C-Terminus (Chain B)
N-Terminus (Chain A)
AraC-LOV Dimer
LBD/DI
Chain A
Chain B

**Supplementary Figure 21: AlphaFold3-predicted structures and associated confidence metrics. a–e,** AlphaFold3 models colored by per-residue confidence (pLDDT), using the following ranges: very high (>90, dark blue), confident (70–90, light blue), low (50–70, yellow), and very low (<50, orange). Each panel is also accompanied by their corresponding predicted aligned error (PAE) matrices. The pLDDT and PAE color bars in panel **a** apply to all panels. For each structure, the N- and C-termini are indicated. **a,** AraC WT, **b,** *As*LOV2 with SG/GS linkers. **c,** WT AraC–LOV fusion. **d,** R2–LOV LdelK variant, **e,** WT AraC–LOV homodimer, which has an interface predicted TM-score (ipTM) of 0.43, indicating low-confidence inter-chain placement. Chains A and B are indicated in the PAE matrix, which shows high error between them except at the ligand-binding domain/dimerization interface (LBD/DI), where interface prediction has higher confidence values; this region is marked in the matrix.
